# Mitochondrial pull-outs are a distinct type of dynamic tubulation events regulated by mitochondrial activity

**DOI:** 10.1101/2025.11.25.690431

**Authors:** Anahita Kasmaie, Priya Gatti, Hema Saranya Ilamathi, Sokhna Mai Gueye, Aidan I. Brown, Uri Manor, Marc Germain

## Abstract

Mitochondrial networks undergo continuous remodeling through fusion and fission, processes that are essential for maintaining energy production and cellular homeostasis. Rapid elongation of the tip of mitochondrial tubules, a process termed dynamic tubulation, also participates in mitochondrial network formation. However, these mechanisms alone cannot fully account for the formation of highly interconnected mitochondrial networks that are required for rapid distribution of mitochondria material. Here, we identify a distinct type of dynamic tubulation, mitochondrial pull-outs characterized by the lateral extrusion from pre-existing mitochondrial tubules, as metabolically regulated determinants of mitochondrial network formation. Pull-outs are distinct from the tip elongation form of dynamic tubulation as they are modulated by the mitochondrial dynamins MFN1 and DRP1 and are stimulated by conditions favoring oxidative phosphorylation. Pull-outs also depend on mitochondrial actin polymerization and are required to increase mitochondrial connectivity and respiratory activity. Together, these findings establish mitochondrial pull-out as a metabolically sensitive mechanism that promotes mitochondrial network connectivity and links organelle architecture to cellular energy demands.

## Introduction

Mitochondria are highly dynamic organelles essential for cellular energy production, metabolic regulation, calcium signaling, and apoptosis. Their morphology and distribution are constantly remodeled through the opposing processes of fission and fusion, which play a central role in maintaining mitochondrial function and structural integrity (*1, 2*). Mitochondrial fusion promotes mixing of contents, protecting against localized defects, while fission facilitates biogenesis, mitophagy, and equal mitochondrial segregation during cell division. The balance between these remodeling events controls mitochondrial bioenergetic output, affecting cell fate and adaptation to stress.

Depending on the cell type and metabolic conditions, mitochondria can be short and isolated or form highly connected networks that promote survival (*3–5*) and have been proposed to facilitate the diffusion of mitochondrial content (*6*). Mitochondrial networks are indeed highly responsive to stress and will rapidly reorganise in response to intracellular and extracellular cues (*3–5, 7*). For example, during amino acid starvation, the formation of highly connected and elongated mitochondrial networks contributes to the maintenance of ATP levels (*3, 4*). Importantly, recent evidence suggests that mitochondrial networks are shaped by events beyond fusion and fission of linear mitochondrial tubules. In fact, most fusion events occur on the side of a mitochondrial tubule rather than at its end, creating new mitochondrial junctions that increase network connectivity (*8, 9*).

In parallel, dynamic tubulation, a process leading to the rapid elongation of a mitochondrial tubule, was shown to be required for mitochondria to populate the periphery of the cell (*10, 11*). While the original study focussed on mitochondria tip elongation(*10*), more recent work included events arising from discrete protrusions on the side of mitochondrial tubules as part of dynamic tubulation events(*10*). Nevertheless, these pull-out events (previously reported in yeast (*12*)) were not fully described and their relationship to mitochondrial tip elongation (the original dynamic tubulation definition(*10*)) remained unclear.

In this study, we identify mitochondrial pull-outs as a distinct form of dynamic tubulation required for the formation of highly connected mitochondrial networks. Mitochondrial pull-outs, but not tip elongation, are stimulated by mitochondrial activity, leading to highly connected mitochondrial networks. Furthermore, pull-outs, but not tip elongation, are regulated by mitochondrial dynamins and mitochondrial actin, further demonstrating that these constitute two distinct types of dynamic tubulation events. Finally, we show that glucose starvation-induced mitochondrial connectivity and oxygen consumption require this process, further supporting an important role for pull-outs in metabolic regulation. Together, these findings establish mitochondrial pull-out as a metabolically regulated remodeling mechanism in mammalian mitochondria that promote mitochondrial network connectivity and cellular adaptation.

## Results

Formation of highly connected mitochondrial networks protects cells from a variety of cellular stresses and is associated with improved mitochondrial metabolism (*3–5*). While changes in mitochondrial structure are usually considered to be the result of mitochondrial fission and fusion, other processes play an important role. Mitochondria can rapidly elongate to populate the periphery of the cell by dynamic tubulation (*10*). While the tip elongation form of dynamic tubulation has been characterised(*10, 11*), pull-out events are still poorly understood (Figure 1A). Pull-outs, but not tip elongation, result in the formation of branched mitochondria that are typical of hyperfused networks associated with cell survival and increased bioenergetics. Thus, these two modes of dynamic tubulation affect mitochondrial network topology in distinct manners and could thus have distinct roles and regulation. We thus set out to characterise mitochondrial pull-outs.

**Figure 1.**
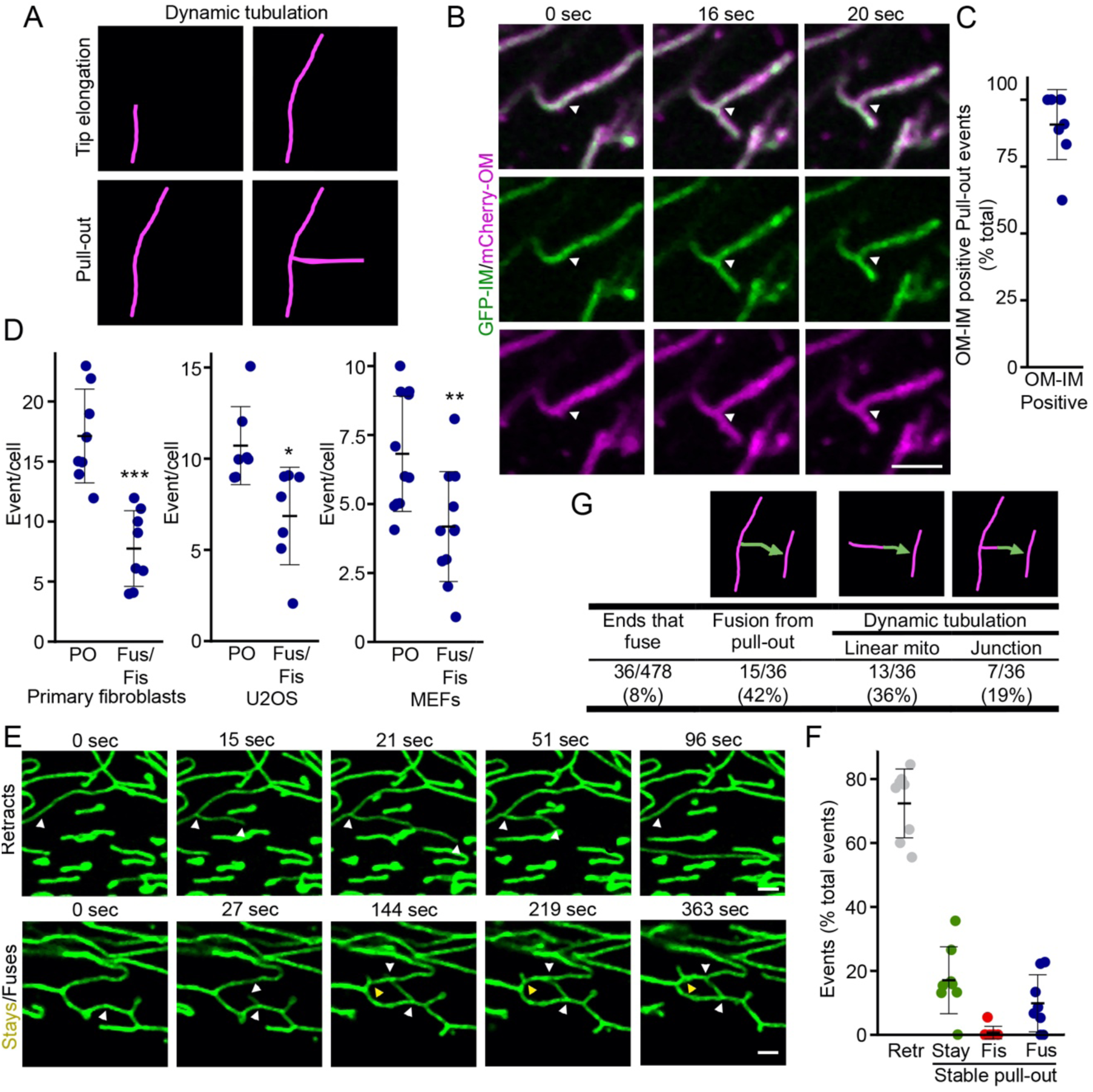
Mitochondrial pull-out events are a frequent feature of mitochondrial networks. (A) Schematic representation of dynamic tubulation events including tip elongation and pull-outs. (B-C) Pull-out events contain outer membrane (mCherry-Fis1, Magenta) and matrix (CCO-GFP, Green) markers. A representative pull-out event in U2OS cells is shown in (B) and quantification in (C). Each point represents an individual cell, with 8 cells quantified in 3 independent experiments. Bars show the average ± SD. (D) Pull-out events are observed in a variety of cell lines (Primary human fibroblasts (Left); U2OS osteosarcoma cells (Middle); Mouse Embryonic Fibroblasts (MEFs) (Right)). The number of pull-out events was plotted with the total number of fusion and fission events from the same video. Each point represents an individual cell, with 8 cells (human fibroblasts, U2OS) and 11 cells (MEFs) quantified in 3 independent experiments. Bars show the average ± SD. Two-sided t-test. * p<0.05, ** p<0.01, *** p<0.001. (E-F) Pull-out events have different fates. (E) Representative images of a retracting (top, white arrowheads), fusing (Bottom, white arrowheads) and stable (Bottom, yellow arrowheads) pull-out event in primary human fibroblasts. (F) Quantification of the number of pull-out events in videos as in (E). Each point represents an individual cell, with 8 cells quantified in 3 independent experiments. Bars show the average ± SD. (G) Pull-out events preferentially fuse. The fate of pull-out events was quantified from 10 cells as in (E-F).

To visualise pull-outs, we transfected U2OS human osteosarcoma cells with outer mitochondria membrane mCherry-Fis1 and matrix-localised CCO-GFP. We readily observed pull-out events in these cells (Figure 1B, Video 1), most of them containing both membranes (Figure 1B-C, Video 2). Pull-outs were also observed in other cell lines (mouse embryonic fibroblasts (MEFs) and primary human fibroblasts, Figure 1D). Interestingly, pull-outs were significantly more frequent than fission and fusion (Figure 1D), suggesting that they provide an important contribution to the shaping of mitochondrial networks.

We observed distinct categories of pull-out behavior across time-lapse recordings of primary fibroblasts stained with mitotracker. Most pull-outs retracted shortly after their formation (Figure 1E (Top), F; mean duration 20±8 seconds, Video 2), representing transient or unstable events. A second group of pull-outs remained for much longer (Supplemental Figure 1A), generating stable mitochondrial branches (Figure 1F (Bottom), Video 3) that could extend several µm (Supplemental Figure 1B; average length 2.8±1.9 µm). Among these stable branches, some successfully fused with other mitochondria (Figure 1F (Bottom, white arrows), G). In rare cases, pull-out events were followed by fission at or near the pull-out site (Figure 1G). In contrast, tip elongation usually lasted longer than pull-outs (Supplemental Figure 1C) but reached similar length (Supplemental Figure 1D).

Interestingly, most fusion events originated from a dynamic tubulation event. In fact, a large fraction of fusion events (42%) involved the end of a mitochondrial tubule generated from a pull-out event, with the rest almost exclusively resulting from tip elongation of a linear or a branched mitochondrion (the latter possibly arising from a previous pull-out)(Figure 1G). As a comparison, only 8% of all mitochondrial ends fused during our imaging period (5 min, Figure 1G). These results indicate that the mitochondrial ends generated by dynamic tubulation, including pull-outs, are selectively used for future fusion events. Overall, our results suggest that mitochondrial pull-outs play an important role in shaping mitochondrial networks.

## Mitochondrial pull-outs and fusion are stimulated by metabolic changes

Mitochondria respond to metabolic changes by altering their length and connectivity (*3, 4, 13, 14*). Therefore, we then asked whether dynamic tubulation participates in these changes in response to changes in nutrient availability. As a model, we first used MEFs grown in galactose instead of glucose, which forces cells to rely on mitochondrial oxidative metabolism. This increased mitochondrial connectivity (Junctions/Ends (*13*)) and total mitochondrial branches, but not mitochondrial length (Figure 2A-B), consistent with previous reports (*3, 4*).

**Figure 2.**
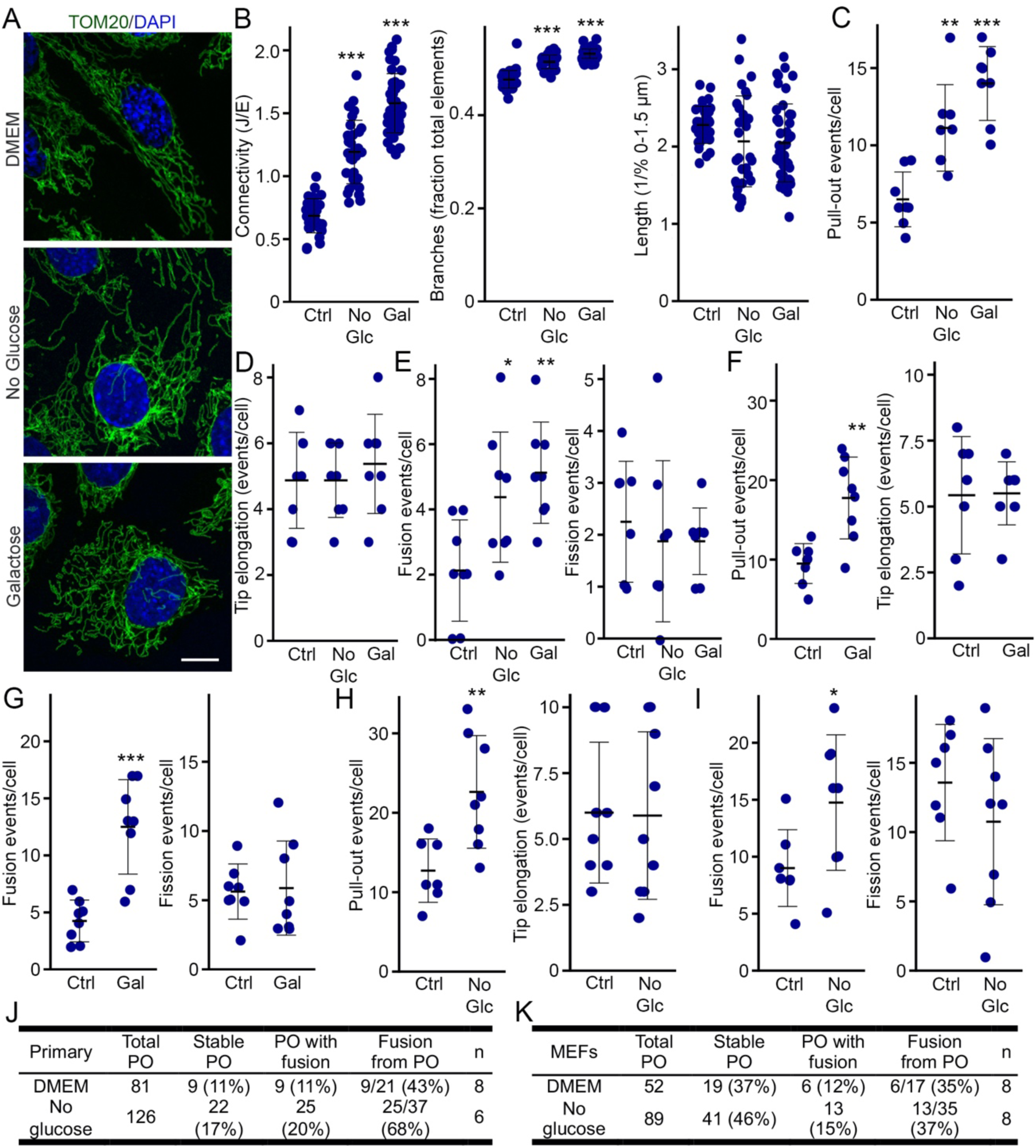
Growth in galactose and glucose starvation promote pull-out events and mitochondrial connectivity. (A-B) Glucose starvation and growth in galactose promote mitochondrial connectivity. (A) Representative images of mitochondrial networks (TOM20, Green; with nuclei marked with DAPI, Blue) in MEFs grown in the indicated conditions. (B) Mitochondrial connectivity (Left), the number of mitochondrial tubules (Middle) and mitochondrial length (Right) were quantified from these images using Momito. Each point represents an individual cell, with 34 (DMEM), 50 (galactose) and 32 (no glucose (Glc)) cells quantified in 3 independent experiments. Bars show the average ± SD. One-way ANOVA. *** p<0.001. (C-D) Quantification of dynamic tubulation under different growth conditions. MEFs were grown for 4 hours in the indicated condition, their mitochondria marked with mitotracker and imaged live. Mitochondrial pull-out (C) and tip elongation (D) were then quantified from the resulting videos. Each point represents an individual cell, with 8 cells per condition quantified in 6 independent experiments. Bars show the average ± SD. Two-sided t-test. ** p<0.01, *** p<0.001. (E) Quantification of mitochondrial dynamics under different growth conditions. Mitohcondrial fusion (Left) and fission (Right) were quantified from the same videos as in (C-D). Each point represents an individual cell, with 8 cells per condition quantified in 6 independent experiments. Bars show the average ± SD. Two-sided t-test. * p<0.05, ** p<0.01, *** p<0.001. (F) Dynamic tubulation in primary human fibroblasts grown in galactose. Pull-outs (Left) and tip elongation (Right) were analysed as in MEFs above. (G) Mitochondrial dynamics in primary human fibroblasts grown in galactose. Fusion (Left) and Fission (Right) were analysed from the same videos as in (F). Each point represents an individual cell, with 8 cells per condition quantified in 6 independent experiments. Bars show the average ± SD. One-way ANOVS. * p<0.05, ** p<0.01, *** p<0.001. (H) Dynamic tubulation in primary human fibroblasts grown in the absence of glucose. Pull-outs (Left) and tip elongation (Right) were analysed as above. (I) Mitochondrial dynamics in primary human fibroblasts grown in the absence of glucose. Fusion (Left) and Fission (Right) were analysed from the same videos as in (H). Each point represents an individual cell, with 8 cells per condition quantified in 6 independent experiments. Bars show the average ± SD. One-way ANOVA. * p<0.05, ** p<0.01, *** p<0.001. (J-K) The fraction of pull-out events leading to fusion is increased in glucose starvation. The number and fate of pull-out events was quantified in primary human fibroblasts (J) and MEFs (K).

Mitochondrial pull-outs were increased under these conditions (Figure 2C), consistent with mitochondrial pull-outs creating new mitochondrial junctions and branches. In contrast, tip elongation was not altered by growth in galactose (Figure 2D), suggesting that tip elongation and pull-outs are regulated in a distinct fashion. Interestingly, MEFs grown in galactose also had an increased number of fusion events, but showed no difference in fission (Figure 2E). We also observed a similar increase in pull-outs and fusion in galactose-treated primary fibroblasts, while fission and tip elongation remained constant (Figure 2F-G). As most fusion events generate new junctions by occurring in a tip-to-side manner (*8, 15*), this suggests that increasing mitochondrial activity mainly promotes the connectivity of mitochondrial networks.

We then assessed the effect of removing glucose from the media. Similar to cells grown in galactose, MEFs and primary fibroblasts subjected to glucose starvation for 4 hours exhibited a marked increase in mitochondrial fusion and a significant elevation in mitochondrial pull-outs (Figure 2C, E, H-I). On the other hand, fission and tip elongation frequency remained unchanged (Figure 2D-E, H-I), further indicating that these events are regulated in a manner that is distinct from pull-outs and fusion.

In primary fibroblasts, there was also an increase in the fraction of pull-outs that were stable or fused, and a greater fraction of the total number of fusion events could be traced to a pull-outs (Figure 2J). The distribution of the increase in pull-outs was somewhat different in MEFs, as both fusion and pull-out events increased in a similar manner upon glucose-starvation and a larger fraction of the pull-outs were stable even in control cells (Figure 2K). Nevertheless, our findings demonstrate that mitochondrial pull-outs and fusion are selectively enhanced under conditions that elevate oxidative metabolism.

## Mitochondrial pull-outs are coupled to mitochondrial activity

Mitochondrial pull-outs, but not tip elongation, are increased in conditions that promote mitochondrial OXPHOS. To directly test the functional link between pull-outs and mitochondrial activity, we used glucose-deprived primary fibroblasts. These cells switch their carbon source to ß-oxidation of lipids upon glucose withdrawal (*16*), as evidenced by a marked reduction in lipid droplets relative to cells grown in the presence of glucose (Figure 3A-B) that was prevented by the ß-oxidation inhibitor etomoxir (Figure 3A-B). Etomoxir also caused a strong decrease in total ATP in the absence of glucose (Figure 3C). This switch to mitochondria-dependent OXPHOS in glucose-starved cells was further demonstrated by the observation that ATP levels were more sensitive to the ATP synthase inhibitor oligomycin in glucose-starved cells relative to control cells (Figure 3C).

**Figure 3.**
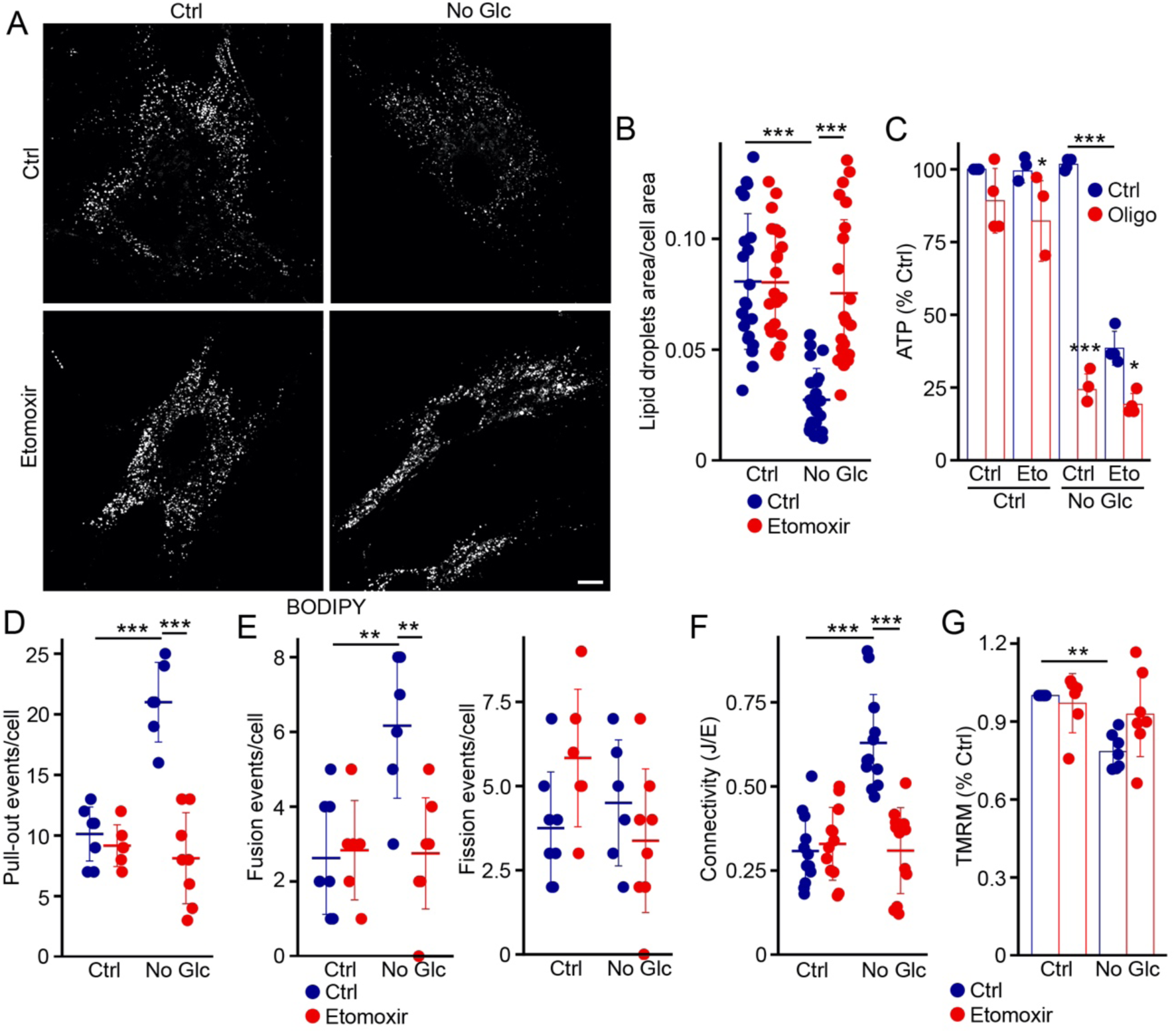
Mitochondrial activity stimulates mitochondrial pull-out. (A-C) Glucose starvation stimulates mitochondrial activity. Primary human fibroblasts were grown for 4 hours in the presence or the absence of glucose with the ß-oxidation inhibitor etomoxir and the ATP synthase inhibitor oligomycin where indicated. Lipid droplets (marked with BODIPY, (A-B) and total ATP levels (C) were then measured. Each point in (B) represents an individual cell, with 22 cells quantified in 3 independent experiments. Each point in (C) represents an experiment. Bars show the average ± SD. One-way ANOVA. * p<0.05, *** p<0.001 (D-E) Quantification of mitochondrial dynamics in response to etomoxir. Primary human fibroblasts were grown for 4 hours in the indicated condition, their mitochondria marked with mitotracker and imaged live. Mitochondrial pull-out (D), fusion (E, Left) and fission (E, Right) were then quantified from the resulting videos. Each point represents an individual cell, with 8 (Ctrl, no glucose with etomoxir) and 6 (etomoxir, no glucose) cells quantified in 3 independent experiments. Bars show the average ± SD. One-way ANOVA. ** p<0.01, *** p<0.001 (F) Etomoxir prevents the increase in mitochondrial connectivity caused by glucose starvation. Cells were grown for 4 hours in the indicated condition, fixed, marked with TOM20, imaged and the resulting images analysed with Momito. Each point represents an individual cell, with 10 cells quantified in 3 independent experiments. Bars show the average ± SD. One-way ANOVA. ** p<0.01, *** p<0.001 (G) Mitochondrial potential in glucose-starved cells. Cells were treated as indicated and membrane potential measured using TMRM. Each point represents an individual experiment, with 7 experiments in total. Bars show the average ± SD. One-way ANOVA. **

Similarly, mitochondrial pull-outs and fusion events were significantly reduced in etomoxir-treated primary human fibroblasts during glucose starvation (Figure 3D-E), indicating that both events are regulated by mitochondrial activity. However, etomoxir did not affect fission (Figure 3E), consistent with the fission protein DRP1 being regulated from the cytosol (*3, 4*). Etomoxir also prevented the increase in connectivity caused by glucose starvation (Figure 3F), indicating that the changes in mitochondrial networks observed under these conditions require mitochondrial pull-outs and fusion. Importantly, the loss of pull-outs and fusion were not the consequence of etomoxir lowering mitochondrial potential, as measured using TMRM (Figure 3G). Overall, our data indicates that mitochondrial activity promotes mitochondrial pull-outs and fusion to reorganize the structure of mitochondrial networks in response to changing metabolic conditions.

## Mitochondrial connectivity promotes diffusion across mitochondrial networks

Mitochondrial elongation in response to amino acid starvation has previously been associated with the maintenance of ATP levels and cristae reorganisation (*3, 14*). In contrast, the changes in carbon source used here mainly alter mitochondrial connectivity, not length, which we postulate improves the flow of metabolites throughout mitochondrial networks to improve their distribution throughout the cell. Consistent with this, these treatments also increased overall mitochondrial area (Figure 4A) but not the expression of mitochondrial proteins, indicating that mitochondrial mass did not change (Figure 4B), suggesting the presence of a larger network that could allow more efficient delivery of metabolites across the cell.

**Figure 4.**
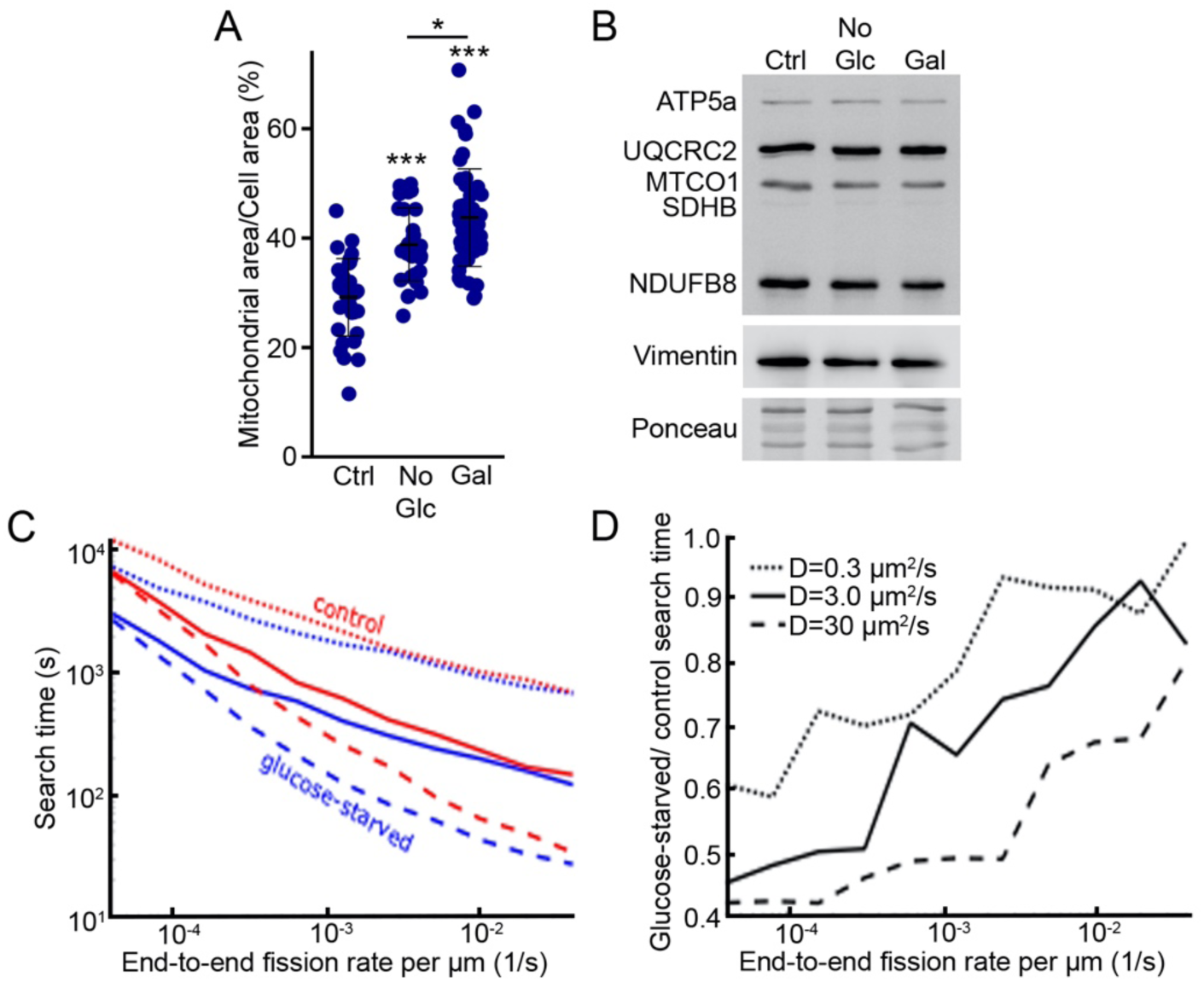
Changes to mitochondrial network structure and dynamics in different metabolic states substantially affects particle spread on the network. (A) Quantification of mitochondrial area (using the mitochondrial marker TOM20) normalised to total cell area in MEFs treated as indicated. Each point represents an individual cell, with 34 (DMEM), 50 (galactose) and 32 (no glucose) cells quantified in 3 independent experiments. Bars show the average ± SD. One-way ANOVA. * p<0.05, *** p<0.001. (B) Western blot showing the expression levels of ETC components in the indicated growth conditions in MEFs. Vimentin and ponceau were used as loading controls. (C) Search time to find a randomly selected mitochondrial fragment from an initial location on another randomly selected fragment as the end-to-end fission rate is varied. Blue curves are for mitochondrial networks with glucose-starved parameters and red curves are for control parameters. Search times averaged over 1000 samples. (D) Ratio of search time on a mitochondrial network with glucose-starved parameter to a network with control parameters, with same data as (C). Across both panels, dotted curves are a particle of diffusivity 0.3 μm^2^/s, solid curves diffusivity 3 μm^2^/s, and dashed curves diffusivity 30 μm^2^/s, as indicated by the legend in (D). Physiological end-to-end fission rate estimated in range 8×10^-4^ to 2.4×10^-3^ events per second per μm (see methods for details).

To address this, we applied a mathematical model of mitochondrial networks combining mitochondrial dynamics with particle diffusion on the network (*6*). To minimize the number of variables, the mitochondrial dynamics parameters were set relative to the end-to-end fission rate using our experimental data (see methods). We then assessed diffusion through the network by calculating the time for a particle to find a randomly selected mitochondrial network location (search time). We found that fast diffusing particles consistent with soluble matrix content (diffusivity of 3 – 30 μm^2^/s) have a substantial 30 – 50% decrease in search time on the more connected networks representing glucose-starved cells compared to control networks (Figure 4C-D). For slower diffusing particles representing membrane proteins (diffusivity of 0.3 μm^2^/s) the decrease in search time is modest, approximately 10%, as the trajectories of slower particles are less affected by connectivity changes (Figure 4C-D). Overall, our model is consistent with mitochondrial connectivity promoting the diffusion of metabolites through the mitochondrial network and the cell to optimise nutrient utilisation.

## Mitochondrial dynamins are associated with pull-outs

Mitochondrial remodeling processes, including fission, fusion and tip elongation occur at ERMCS. We thus determined if this was also the case for mitochondrial pull-outs. For this, we performed live-cell imaging of HeLa cells stably expressing ER-mTurquoise in which mitochondria were visualised with mitotracker (Figure 5A, Video 4). Strikingly, 95% of pull-out events were localized at ER-mitochondria interfaces (Figure 5B), suggesting that these events occur at ERMCS.

**Figure 5.**
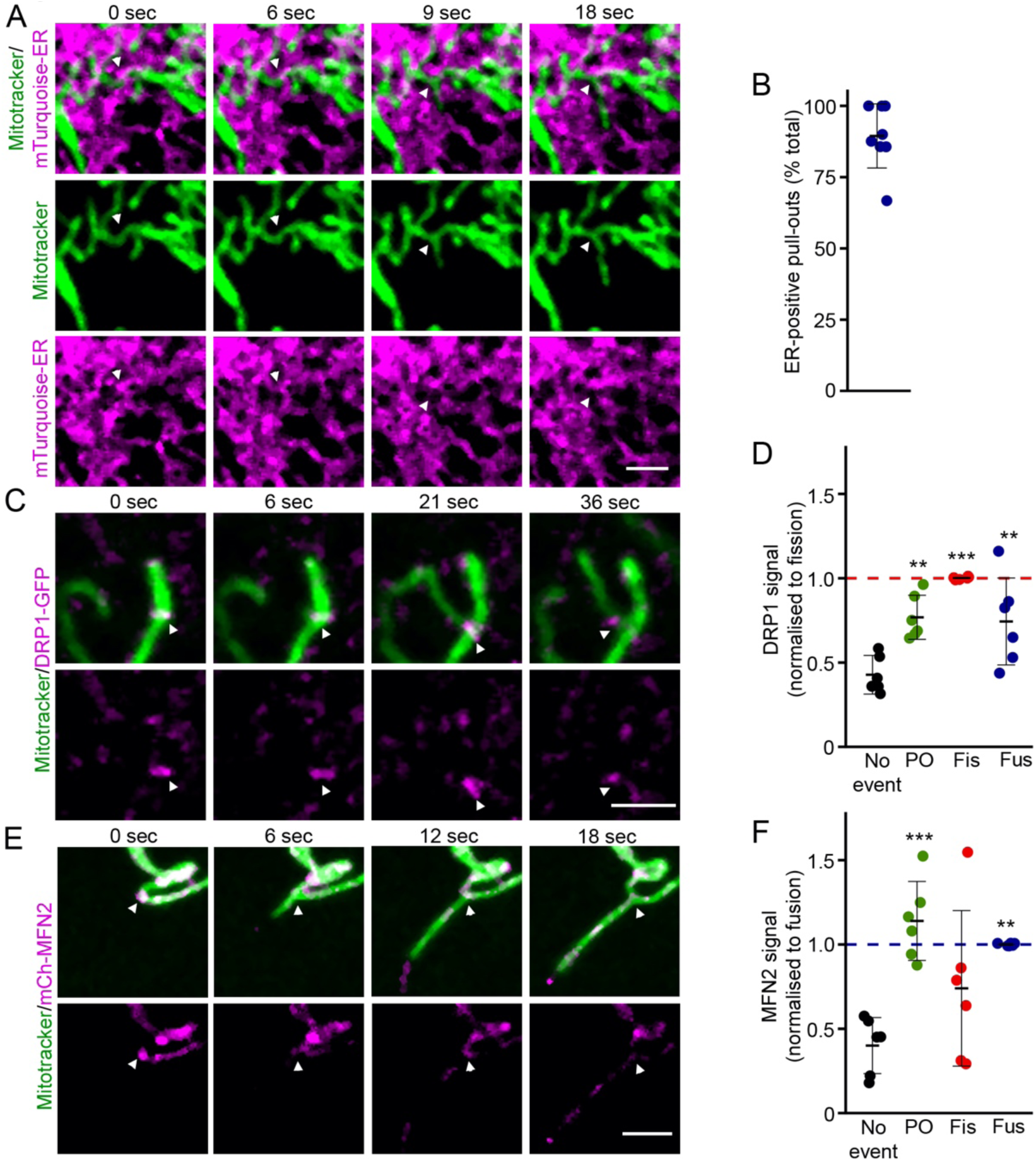
Pull-out events occur at mitochondrial fission and fusion hotspots. (A-B) The ER is present at pull-out events. (A) Representative images from HeLa cells stably expressing mTurquoise-ER (magenta) and imaged live with mitochondria marked with mitotracker (Green). Arrowheads denote a pull-out event. Scale bar 2 µm. (B) Quantification of the pull-out events positive for mTurquoise-ER. Each point represents an individual cell, with 9 cells quantified in 3 independent experiments. (C-D) Recruitment of DRP1 at pull-out sites. (C) representative images of cells transfected with DRP1-GFP (Magenta) and imaged live in the presence of mitotracker to mark mitochondria. Arrowheads denote a pull-out event. Scale bar 2 µm. (D) Quantification of the enrichment of DRP1-GFP at different types of events. The data was normalised to the average DRP1-GFP signal at fission sites for each cell. Each point represents an individual cell, with 6 cells quantified in 3 independent experiments. Bars show the average ± SD. One-way ANOVA. ** p<0.01, *** p<0.001. (E-F) Recruitment of MFN2 at pull-out sites. (E) representative images of cells transfected with mCh-MFN2 (Magenta) and imaged live in the presence of mitotracker to mark mitochondria. Arrowheads denote a pull-out event. Scale bar 2 µm. (F) Quantification of the enrichment of mCh-MFN2 at different types of events. The data was normalised to the average mCh-MFN2 signal at fusion sites for each cell. Each point represents an individual cell, with 6 cells quantified in 3 independent experiments. Bars show the average ± SD. One-way ANOVA. ** p<0.01, *** p<0.001.

During fission and fusion, ERMCS act as hotspots enriched in actin, where the fission and fusion dynamins (DRP1 and MFN1/2) are recruited(*8, 15*). However, dynamic tubulation was previously shown to be independent of the fission protein DRP1 and the fusion proteins MFN1/2(*10*). To determine if this applied to both tip elongation and pull-outs, we first measured the recruitment of the fission protein DRP1 and the fusion protein MFN2 to pull-out sites since both proteins converge to mitochondria hotspots before fission or fusion occurs (*8, 15*). For this, we first transfected primary human fibroblasts with a GFP-tagged version of DRP1 and imaged them live in the presence of mitotracker to mark mitochondria. DRP1 is known to make foci along mitochondria that colocalize with sites of mitochondrial fission and fusion (*8, 15*). Consistent with this, DRP1 was also present at the base of pull-out events (Figure 5C, Video 5). This accumulation of DRP1 at the base of pull-out events was quantified by measuring the enrichment of the fluorescent signal at pull-out sites relative to the fluorescence signal at fission sites within the same cell. While the enrichment of DRP1 at pull-out sites was somewhat less than at fission events, its enrichment was still significantly above the DRP1 signal present at sites where no event was occurring (Figure 5D).

We then repeated the experiment in the presence of mCherry-MFN2 (mCh-MFN2) instead of DRP1. Consistent with mitochondrial hotspots being the site where pull-outs originates, MFN2 was present at pull-out sites where its signal was enriched to a similar extent as fusion events (Figure 5E-F, Video 6). Overall, our results indicate that pull-out events occur at mitochondrial hotspots, similar to fusion and fission.

As pull-out events occur at mitochondrial dynamics hotspots enriched in both fission and fusion dynamins, we then determined whether these proteins play a role in mitochondrial pull-out. For these experiments, we used short-term transient knockdown of DRP1 or MFN1 to minimise mitochondrial defects that could affect the analysis (Figure 6A-C). DRP1 knockdown resulted in highly elongated mitochondria (Figure 6B, D), but did not alter overall mitochondrial connectivity (Figure 6D). This is the result of a fraction of the mitochondrial network that was more connected in the perinuclear area (Figure 6B), consistent with this area later developing mitobulbs (*17, 18*). As expected(*10*), loss of DRP1 did not significantly affect tip elongation (Figure 6E), validating that mitochondria are still dynamic in the knockdown conditions we used. However, pull-out events were strongly reduced in these cells (Figure 6E), suggesting that, contrary to tip elongation, DRP1 participates in pull-outs. To further ensure that the observed differences were not the result of altered mitochondrial networks, we normalised tip elongation and pull-out events to mitochondrial size and found very similar results (Figure 6F).

**Figure 6.**
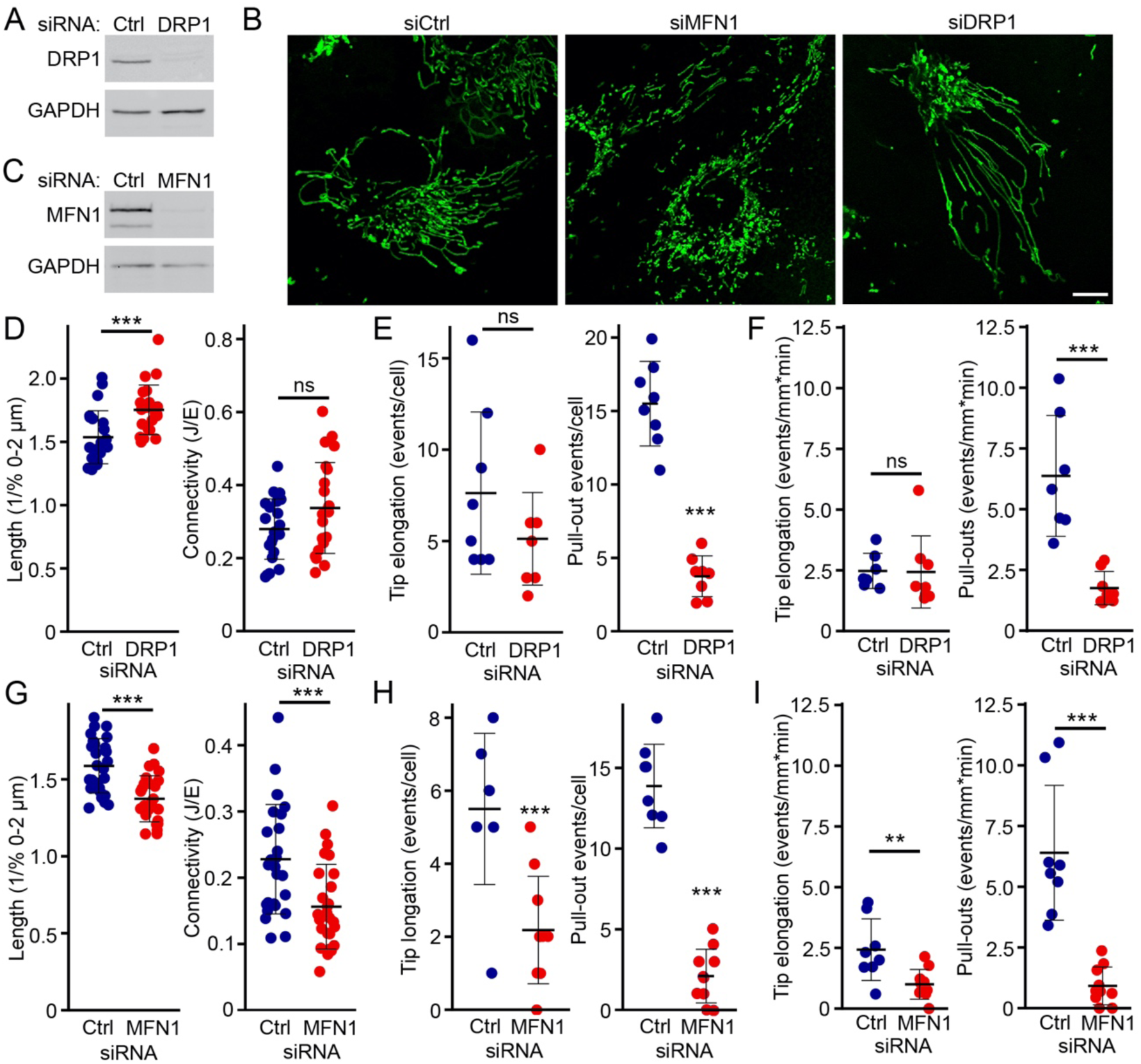
MFN1 and DRP1 are required for mitochondrial pull-out. (A-C) Knockdown of DRP1 or MFN1 affect mitochondrial structure. (A) Expression of DRP1 in control and siDRP1-treated U2OS cells. GAPDH was used as a loading control. (B) Representative images from U2OS cells in which DRP1 or MFN1 has been knocked down. Mitochondria were marked with an antibody against TOM20. Scale bar 10 µm. (C) Expression of MFN1 in control and siMFN1-treated U2OS cells. GAPDH was used as a loading control. (D-F) Knockdown of DRP1 prevents mitochondrial pull-out. (D) Quantification of mitochondrial length (Left) and connectivity (Right) from images as in (B) control and siDRP1-treated U2OS cells. Each point represents an individual cell, with 22 cells quantified in 3 independent experiments. Bars show the average ± SD. Two-sided t-test. *** p<0.001. (E-F) Quantification of tip elongation (Left) and pull-outs (Right) from live cell imaging of U2OS transfected with the indicated siRNA as total number of event/5 minute videos (E) or normalised to total mitochondrial length and time (F). Each point represents an individual cell, with 7 cells quantified in 3 independent experiments. Bars show the average ± SD. Two-sided t-test. *** p<0.001. (G-I) Knockdown of MFN1 prevents mitochondrial pull-out. (G) Quantification of mitochondrial length (Left) and connectivity (Right) from images as in (B) control and siMFN1-treated U2OS cells. Each point represents an individual cell, with 26 cells quantified in 3 independent experiments. Bars show the average ± SD. Two-sided t-test. *** p<0.001. (H) Quantification of tip elongation (Left) and pull-outs (Right) from live cell imaging of U2OS transfected with the indicated siRNA as total number of event/5 minute videos (H) or normalised to total mitochondrial length and time (I). Each point represents an individual cell, with 8 cells quantified in 3 independent experiments. Bars show the average ± SD. Two-sided t-test. *** p<0.001.

We then used a similar approach to assess the role of MFNs. Since MFN2 has several roles while MFN1 in mainly involved in fusion (*19*), we specifically addressed the role of MFN1. Knockdown of MFN1 resulted in mitochondria that were shorter and less connected, but not fragmented (Figure 6B, G). Consistent with their lower connectivity, siMFN1 cells showed strongly reduced pull-out frequency both in total number and relative to network size (Figure 6H-I), suggesting a critical role for MFN1 in the initiation of pull-out events. Interestingly, and in contrast to what was previously reported(*10*), tip elongation was also affected in siMFN1 cells, although to a lesser degree than pull-outs (Figure 6H-I). Overall, our results indicate that tip elongation and pull-outs are different types of dynamic tubulation regulated in a distinct manner, both in terms of metabolic regulation and dependence on mitochondrial dynamins.

## Actin regulates pull-outs in response to metabolic changes

Actin polymerisation at fission and fusion sites plays a key role in the control of mitochondrial dynamics (*8, 20, 21*). To determine whether pull-out events similarly require actin, we first transfected primary fibroblasts with mitochondria-targeted actin chromobodies (AC-mito)(*8, 21*) to visualise mitochondrial actin, along with mCherry-Fis1 to label mitochondria. Most pull-out sites were marked by AC-mito signal, which also extended into the newly formed mitochondrial branch (Figure 7A, quantified in 7B). As these results indicated that actin is present at the pull-out site, we then determined its role.

**Figure 7.**
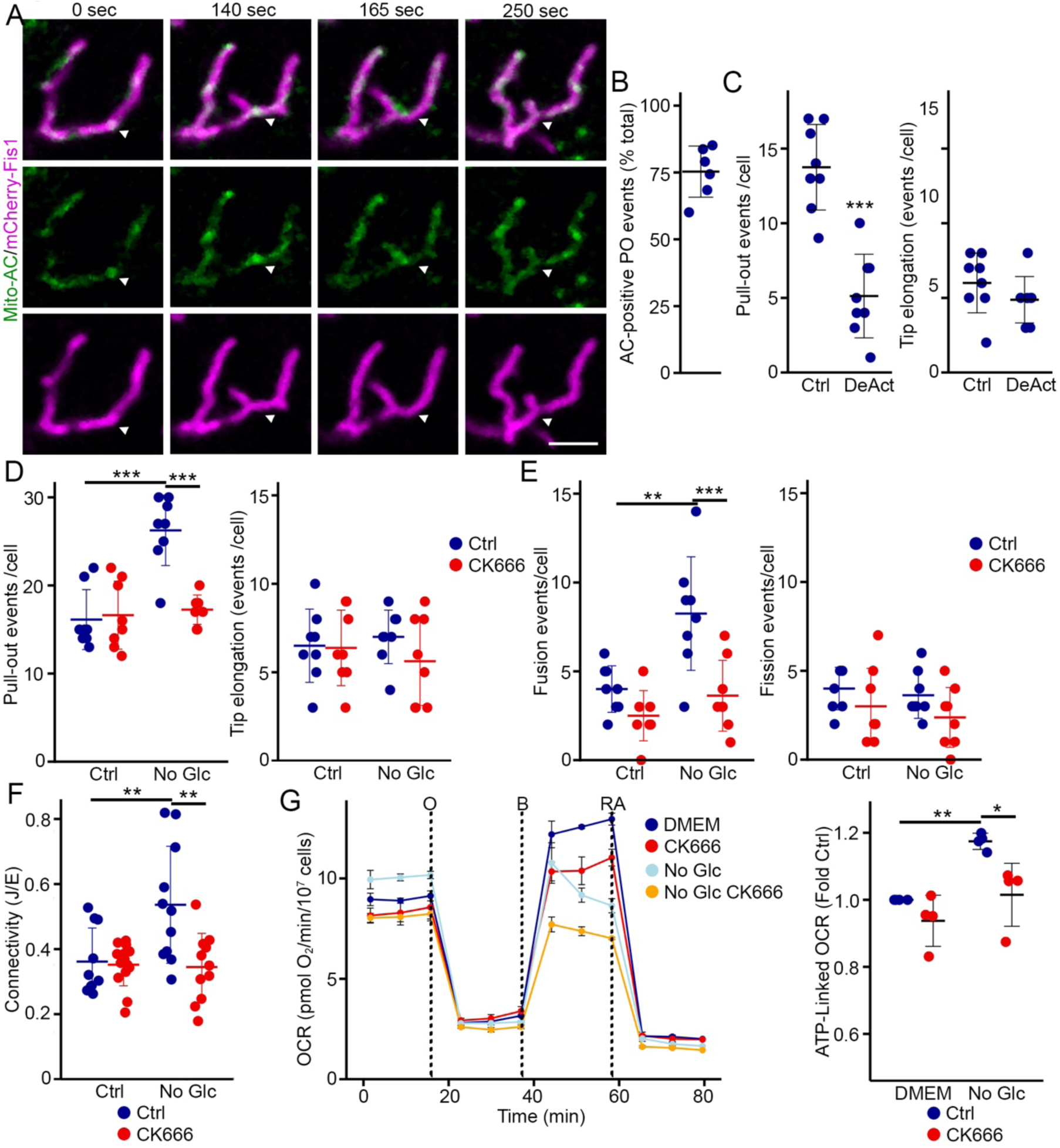
Actin regulates pull-out events in glucose-starved cells. (A-B) Actin is present at pull-out sites. (A) Representative pull-out event showing the enrichment of mitochondrial actin. Cells were transfected with a GFP-tagged mitochondria-targeted actin chromobody (mitoAC, Green) and mitochondria-targeted mCherry (mCherry-Fis1, Magenta) and imaged live. The arrowhead denotes the pull-out event. Scale bar 2 µm. (B) Quantification of pull-out events positive for AC-mito. Each point represents an individual cell, with 6 cells quantified in 3 independent experiments. Bars show the average ± SD. (C) Quantification of pull-outs (Left) tip elongation (Right) in cells transfected with DeAct-mito. Each point represents an individual cell, with 8 cells quantified in 3 independent experiments. Bars show the average ± SD. Tow-sided t-test. *** p<0.001. (D-E) Quantification of pull-outs (D, Left) tip elongation (D, Right), as well as fusion (E, Left) and fission (E, Right) in cells treated with the Arp2/3 inhibitor CK666. Each point represents an individual cell, with 8 cells quantified in 3 independent experiments. Bars show the average ± SD. Two-way ANOVA. ** p<0.01, *** p<0.001. (F) CK666 prevents the increase in connectivity caused by glucose starvation. Cells were glucose starved for 4 hours before mitochondrial networks were analysed. Each point represents an individual cell, with at least 10 cells quantified in 3 independent experiments. Bars show the average ± SD. Two-way ANOVA. ** p<0.01. (G) CK666 prevents the increased in oxygen consumption (OCR) caused by glucose starvation. (Left) Representative experiment with 6 replicates, Bars show the average ± SD. O, Oligomycin; B, Bam15; RA, Rotenone and Antimycin. (Right) Quantification of ATP-linked OCR. Each point represents an experiment. Bars show the average ± SD. Two-way ANOVA. * p<0.05, ** p<0.01.

We first selectively removed actin from mitochondria using the mitochondria-targeted DeAct tool (DeAct-mito) we previously published (*8*). U2OS cells were transfected with DeAct-mito and dynamic tubulation was scored in from live cell imaging. Consistent with the presence of actin at pull-out sites, DeAct-mito prevented pull-out formation (Figure 7C). In contrast, tip elongation was not affected (Figure 7C), further supporting the idea that these are two independent types of dynamic tubulation.

We previously showed that the branch actin nucleation complex Arp2/3 is required for fusion but not fission (*8*). As pull-outs and fusion are induced by glucose starvation, we then addressed the role of the Arp2/3 complex in this process. For this, we inhibited the Arp2/3 complex with CK666 and determined the effect on mitochondrial dynamics. Consistent with a role for actin in mitochondrial pull-outs, CK666 prevented the increase in pull-outs caused by glucose starvation (Figure 7D), but did not affect tip elongation (Figure 7D). CK666 also inhibited fusion but not fission (Figure 7E), consistent with our previous report (*8*). Consistent with this, Arp2/3 inhibition prevented the increase in mitochondrial connectivity caused by glucose starvation (Figure 7F), indicating an important role for actin in the control of mitochondrial structure under changing metabolic conditions.

We then determined the impact CK666 on mitochondrial oxygen consumption (OCR). Consistent with glucose-starved cells being more dependent on their mitochondria for ATP production, ATP-linked OCR was increased in these cells (Figure 7G). Importantly, CK666 prevented the increase in OCR caused by glucose starvation, indicating that actin-dependent pull-outs and fusion are required for cells to adapt their mitochondrial output to variations in nutrient availability. Overall, our work indicates that the two forms of dynamic tubulation that were previously reported, tip elongation and pull-outs, are regulated in a distinct manner and that the increase in pull-outs caused by glucose starvation is required for cells to adapt their mitochondrial output.

## Discussion

Mitochondrial dynamics play a key role in the maintenance of mitochondrial morphology, bioenergetics, distribution, and quality control. Mitochondrial fission and fusion are generally thought to be the main processes balancing mitochondrial fragmentation and elongation to match cellular metabolic and signaling demands (*1, 2*). However, these processes do not fully explain the diversity of mitochondrial networks, especially with regards to their connectivity. Here, we show that mitochondrial pull-out is a distinct type of mitochondrial behaviour required for the remodelling of mitochondrial networks occurring in response to metabolic alterations.

Mitochondria change their length and connectivity in response to different cellular cues. The generation of hyperfused mitochondrial networks was first reported as a response to mild cellular stress (*5*) and amino acid starvation (*3, 4*). The mitochondrial elongation caused by starvation was further shown to depend on the inhibition of mitochondrial fission (*3*), although this did not explain why mitochondrial networks also become more connected under these conditions (*13*). In this context, recent research has expanded our view of mitochondrial dynamics by highlighting the intricate nature of fission and fusion and introducing additional remodeling processes (*1*). For example, we now know that there are two distinct types of fission (tip and mid-zone (*22*)) and that most fusion events occur with the tip of a mitochondrion fusing with the side of another one, creating a new junction (*8, 9*). In addition, dynamic tubulation supports intracellular distribution of mitochondria and spatial plasticity (*10*). However, the role of these events in shaping the response of mitochondrial networks to metabolic challenges has remained poorly understood.

We show here that metabolic alterations promoting mitochondrial activity stimulate the formation of highly connected mitochondrial networks through the activation of mitochondrial pull-out and fusion. Pull-out events generate mitochondrial tips that are especially prone to fusion, allowing a rapid increase in mitochondrial connectivity that increases mitochondrial area. It is noteworthy that these events occur without changes in mitochondrial fission or tip elongation. This is consistent with the previous observation that amino acid starvation, but not glucose deprivation, triggers mitochondrial elongation through the inhibition of mitochondrial fission (*3, 4*).

The increase in mitochondrial connectivity and area observed in response to increased pull-out events likely allows a better usage of mitochondrial resources, either by allowing a better access to nutrients or by promoting the distribution of the resulting ATP. Consistent with this, we found that connected networks modeled from glucose-starved cells allowed faster distribution of fast-diffusing soluble content than their equivalent from glucose-fed cells. This is consistent with more connected networks being more efficient at spreading their content (*6*). This also suggests that promoting pull-outs and fusion (increasing connectivity) will have an effect on network configuration and diffusion that is distinct from inhibiting fission, which is more associated with mitochondrial elongation (*3, 4*). Indeed, inhibition of Arp2/3-dependent pull-outs and fusion blocked the increase in OCR caused by glucose starvation.

Dynamic tubulation was originally defined as thin, motile tubules extending along microtubules (*10*), which we defined here as tip elongation to distinguish it from pull-outs that originate from the side of a mitochondrial tubules. While these two forms of dynamic tubulation were previously considered as the same process (*10*), our results demonstrate that they represent mechanistically and functionally distinct processes. First, pull-outs, but not tip elongation, are increased in response to glucose starvation and growth in galactose. Second, contrary to tip elongation, pull-outs require actin and the presence of mitochondrial dynamins.

While mitochondrial dynamins are required for pull-out events, the underlying signalling mechanisms remain elusive. Given that pull-out events are stimulated by mitochondrial activity, one possibility is that the signal is provided by a mitochondrial metabolite. This would be similar to the regulation of cristae structure by the fusion protein OPA1 that is controlled in response to metabolic changes through its interaction with members of the SLC25A family of mitochondrial solute carriers (*14*).

In summary, our findings identify mitochondrial pull-out as a novel and metabolically regulated remodeling mechanism that complements and expands the classical model of mitochondrial dynamics. Pull-out events generate fusion-competent ends that actively increase network connectivity. By leveraging both fusion and fission machinery but responding directly to metabolic demand, pull-out integrates structural, spatial, and energetic remodeling into a unified adaptive process. This would allow faster diffusion of matrix components and optimization of metabolite distribution. This discovery enriches the conceptual landscape of mitochondrial dynamics, opening new avenues for understanding organelle homeostasis and targeting mitochondrial dysfunction in disease. Future studies will define the upstream signals, structural determinants, and physiological consequences of pull-out formation.

## ACKNOWLEDGEMENTS

This work was supported by grants from the Natural Sciences and Engineering Research Council of Canada to MG and AIB and Fonds de Recherche du Québec-Nature et Technologies (#2024-PR-326143) to MG. AIB was also supported by start-up funds from the Toronto Metropolitan University Faculty of Science, and by the Toronto Metropolitan University Faculty of Science Dean’s Research Fund. UM is supported by funds from the Dr. David V. Goeddel Endowed Chair in Biological Sciences, the Goeddel Family Technology Sandbox, Charcot-Marie-Tooth Association, Charcot-Marie-Tooth Research Foundation, and the MFN2 Foundation. AK is a recipient of a CERMO-FC scholarship. PG and HSI were supported by a Queen Elizabeth II Diamond Jubilee Scholarship. PG also received a FRQ-S doctoral scholarship, while HSI received a FRQ-NT doctoral scholarship.

## AUTHOR CONTRIBUTIONS

Conceptualization, AK, PG, HSI, AIB, UM and MG; Methodology, AK, PG, HSI, AIB, UM and MG; Investigation, AK, PG, HSI and MG; Writing–Original Draft, AK and MG; Writing – Review and Editing, all the authors; Supervision, MG; Funding Acquisition, AIB, UM, MG

## Materials and Methods

### Cell Culture

Primary human fibroblasts (purchased from the Coriell institute) were cultured in Dulbecco’s Modified Eagle Medium (DMEM) supplemented with 10% fetal bovine serum (FBS) and 100 IU/mL penicillin/100 µg/mL streptomycin. HeLa cells stably expressing mTurquoise-ER, U2OS and mouse embryonic fibroblasts (MEFs) were cultured under the same conditions. For glucose starvation experiments, cells were incubated in glucose-free DMEM supplemented with 10% FBS, L-glutamine, and sodium pyruvate for 4 hours. For galactose conditions, cells were cultured in glucose-free DMEM containing 4.5 g/L galactose, 10% FBS, L-glutamine, and sodium pyruvate for 4 hours. Where indicated, cells were treated with 20 nM etomoxir (Sigma-Aldrich) for 4 hours or 10 µM CK-666 (CAS 442633-00-3 – Calbiochem) for 1 hour prior to imaging or analysis.

### Plasmids and Transient Transfection of Primary Cells

Primary fibroblasts were transfected as previously described(*23*). Briefly, cells were trypsinized and centrifuged at 5000 rpm for 5 minutes. The pellet was resuspended in 10 µL of Neon transfection buffer R (Thermo Fisher Scientific, MPK1096). Cells were transiently transfected with mCherry-MFN2 (Addgene #141156), GFP-DRP1 (Addgene #191942), mCherry-Fis1 (*21*), AF-Fis1 (*21*) and Cox8A-EGFP (Addgene #134861). For all experiments, 1 µg total DNA was used per 10 µL of 10⁶ cells/mL (either individually or combined for co-transfections). Live-cell imaging or immunofluorescence was performed 24 hours post-transfection.

### siRNA Treatments

U2OS cells were seeded in 6-well dishes to reach 60–70% confluency within 24 hours, then transfected with 10 nM MFN1 siRNA (Dharmacon, siGenome, cat#M-010670-01-0005), DRP1 siRNA (Thermo Fisher Scientific, Silencer Select, cat#4390771), or control siRNA (Thermo Fisher Scientific, Silencer Select, 4390843) using siLenFect lipid reagent (Bio-Rad, 1703361). Cells were imaged live or collected for Western blot analysis 24 hours post-transfection.

### ATP Assay

MEFs (30,000 cells) and primary cells (10,000 cells) were seeded in 96-well black plates. Following treatments, ATP content was assessed using the CellTiter-Glo Luminescent Cell Viability Assay reagent (Promega) according to the manufacturer’s instructions. 50 µl of CellTiter-Glo reagent was added to 100 µl of cell culture media in each black-well plate and incubated for 10 minutes at room temperature to stabilize the luminescent signal. Luminescence was measured using a BioTek microplate reader plate reader. To normalize ATP values to protein content, the same number of cells per condition was seeded in separate transparent 96-well plates. After treatments, cells were lysed in 20 µl lysis buffer (10 mM Tris-HCl, pH 7.9, 150 mM NaCl, 1% Triton X-100, 1 mM EDTA) per well, mixed with 20 µl of DC™ Protein Assay Reagent A (Bio-Rad, Cat. #5000113) plus DC™ Protein Assay Reagent S (Bio-Rad, Cat. #500-0115), incubated for 5 minutes, followed by addition of 150 µl DC™ Protein Assay Reagent B (Bio-Rad, Cat. #5000114), then incubated for 15 minutes at room temperature. Absorbance was measured on BioTek microplate reader according to manufacturer’s protocol.

### Oxygen consumption rates

Oxygen consumption rates (OCR) were measured using the Seahorse XF Cell Mito Stress Test Kit (Agilent, Cat. 103708-100) on a Seahorse XFe96 analyzer (Agilent) according to the manufacturer’s instructions. Cells were seeded at 15,000 cells per well in Seahorse XF96 cell culture plates and incubated for 24 h at 37°C and 10% CO2. For CK666-treated conditions, cells were pretreated with CK666 for 1 h before the assay. The culture medium was then replaced with XF DMEM assay medium supplemented as indicated by the manufacturer, and the plates were incubated for 1 h at 37°C in a non-CO2 incubator before measurement. OCR was recorded following sequential injections of oligomycin (1.5 μM), BAM15 (0.25 μM), and rotenone/antimycin A (0.5 μM). Basal OCR and ATP-linked OCR were determined according to the manufacturer’s instructions.

### Lipid droplets visualisation

Cells were seeded onto coverslips and incubated for 24 hours to allow attachment. Subsequently, 5 µM BODIPY FL C12 HPC (Invitrogen, D3792) was added directly to the culture media and incubated with the cells for 18 hours then treated as indicated. Cells were then fixed with paraformaldehyde and imaged.

### TMRM

Following treatments, cells were incubated with 50 nM TMRM in media for 30 minutes at 37°C, protected from light to prevent photobleaching. Cells were then trypsinized, washed once with PBS and resuspended in 200 µl PBS. TMRM Fluorescence intensity was measured using a Cytoflex FACS analyzer (Beckman) with excitation at 514 nm and detection at 570 nm.

### Immunofluorescence

Cells were seeded on glass coverslips (Fisherbrand, 1254580) and allowed to adhere overnight. Cells were then treated, fixed with 4% paraformaldehyde for 15 minutes at room temperature, permeabilized with 0.2% Triton X-100 in PBS, and blocked with 1% BSA/0.1% Triton X-100 in PBS. Coverslips were then incubated with a TOM20 antibody (Rb, Abcam, ab186735, 1:250), followed by fluorophore-conjugated secondary antibodies (Jackson ImmunoResearch, 1:500) and counterstained with DAPI (Invitrogen, D1306, 1:100).

### Live imaging with confocal microscope

Cells were seeded onto glass-bottom dishes and, where indicated, incubated with 100 nM MitoTracker Deep Red (Thermo Fisher Scientific, M7512) in pre-warmed medium for 5 minutes at 37 °C. After staining, cells were washed three times with phosphate-buffered saline (PBS), then maintained in complete, glucose-free, or galactose-containing medium as indicated for each experiment. Live-cell imaging was performed at 37 °C using a Leica Stellaris 5 confocal microscope (63×/1.40 NA oil objective). Time-lapse images were acquired at 0.33–0.78 frames per second for a total imaging duration of 5 minutes. For three-dimensional (3D) reconstruction, Z-stacks were acquired comprising 5 optical sections at 3-second intervals between frames.

For experiments involving MFN2 or DRP1 manipulations, imaging was conducted on a Zeiss LSM 980 Airyscan confocal microscope equipped with a Plan-Apochromat 63×/1.40 Oil DIC M27 objective. Time-lapse series were acquired at 5–11.5 seconds per frame for 10 minutes, and images were processed for Airyscan super-resolution using Zen Blue software (2D-SR, auto-filter settings).

### Image Processing and Analysis

All image processing was performed in ImageJ/Fiji. Unless otherwise noted, images represent single focal planes. Mitochondrial area was measured by segmenting images using Median filter (1.0), followed by thresholding and binary erosion. Total area was measured using the “Measure” function. For the quantification of lipid droplets, the images were segmented as above and droplet numbers were determined using “Analyze Particles” and normalized to total cell size.

### Tracking of Mitochondrial Pull-Out, Fusion, and Fission Events

Fusion and fission events were manually tracked frame-by-frame. Fission was defined as the stable separation of mitochondrial markers. Fusion events were counted when two mitochondria remained fused for ≥100 seconds (*24*). Pull-out events were defined as the emergence of a new mitochondrial branch from the side of a mitochondrion that either retracted, remained stable, or fused with another mitochondrion. ER colocalization at sites of fission/fusion was assessed manually.

### Quantification of mCherry-MFN2 and GFP-DRP1 Signal Accumulation

Regions of interest (0.8 µm²) were drawn around sites of apparent MFN2 or DRP1 accumulation. Mean pixel intensity of GFP/mCherry channels was measured in these regions and normalized to the average DRP1 (for fission) or MFN2 (fusion and pull-out) intensity for that cell. The values were then averaged per cell.

### Mitochondrial Length and Connectivity

Length and connectivity were analyzed using the Momito algorithm (*13*). Mitochondrial networks were segmented using the Tubeness plugin and default thresholding in Fiji, followed by processing in Momito. Connectivity was calculated as the ratio of junctions (J) to ends (E) as previously described (*13*).

### Western Blot

Cells were lysed in 10 mM Tris-HCl (pH 7.4), 1 mM EDTA, 150 mM NaCl, and 1% Triton X-100, supplemented with protease and phosphatase inhibitor cocktails (Sigma-Aldrich). Lysates were centrifuged at 16,000 x g for 10 minutes, and supernatants were collected. Protein concentration was determined by DC protein assay (Bio-Rad). For SDS-PAGE, 20 µg protein was mixed with 2× Laemmli buffer containing β-mercaptoethanol, separated by SDS-PAGE, transferred to nitrocellulose membranes, and probed with primary antibodies against OXPHOS (Mouse 1:1000, Abcam #ab110413), MFN1 (Rabbit, 1:1000; Abcam #ab221661), DRP1 (Mouse, 1:1000, BD #611112), Vinculin (Rabbit, 1:10,000, Cell Signaling Technology mAb13901), and β-Actin (Mouse, 1:10,000, SCBT sc-47778). HRP-conjugated secondary antibodies (Jackson ImmunoResearch, 1:5000) were used, and bands were visualized using enhanced chemiluminescence (Thermo Fisher Scientific) on a Bio-Rad imaging system.

### Simulations

We simulated mitochondrial dynamics, particle diffusion, and search as previously described (*6*). We used 50 minimal mitochondrial fragments, which would correspond to a large yeast cell (*25*). With 30 minimal fragment previously corresponding to a typical yeast cell (*25*) and a total yeast mitochondrial network length of 75 μm (*25*), each minimal fragment is 2.5 μm long. A rate 1 per second for a particle to move from one minimal fragment to a neighboring connected minimal fragment corresponds to a diffusivity of approximately 3 μm^2^/s.

To estimate fission and fusion parameters, the mitochondrial network is assumed to have reached steady state. This can be expressed mathematically as *b*_1_*N*_2_ = *a*_1_(*N*_1_)^2^, for end-to-end fusion balanced by end-to-end fission; and *b*_2_*N*_3_ = *a*_2_*N*_1_*N*_2_, for end-to-side fusion balanced by end-to-side fission. *a*_1_ is the end-to-end fusion rate constant, *b*_1_ the end-to-end fission rate constant, *a*_2_ the end-to-side fusion rate constant, and *b*_2_ the end-to-side fission rate constant, *N*_1_ the number of ends (degree one nodes), *N*_2_ the number of fused ends (degree two nodes), and *N*_3_ the number of three-way junctions (degree three nodes). *b*_2_ = (3/2)*b*_1_ (*26*). This allows the expression of the fusion rates in terms of the end-to-end fission rate *b*_1_: *a*_1_ = *b*_1_*N*_2_/(*N*_1_)^2^ and *a*_2_ = 3*b*_1_*N*_3_/(2*N*_1_*N*_2_).

We set the model parameters according to our experimental data for control and glucose-starved cells. Our experiments measure *N*_1_ = 0.43*N*_nodes_, *N*_2_ = 0.42*N*_nodes_, and *N*_3_ = 0.14*N*_nodes_ for control cells; and *N*_1_ = 0.36*N*_nodes_, *N*_2_ = 0.45*N*_nodes_, *N*_3_ = 0.18*N*_nodes_. Our experiments also measure a mean of 14.5 end-to-end fission events per cell in five minutes, which is 0.048 fission events per second per cell. Fibroblast mitochondria are approximately 5% of the cell area in images (*27*), which we assume transfers to an equal mitochondrial volume fraction of 5%. Fibroblasts are approximately 1600 μm^3^ (*28*), for which mitochondria have an 80 μm^3^ volume for 5% mitochondrial volume fraction, which for a mitochondrial diameter of approximately 700 nm (*29*) corresponds to approximately 200 μm in mitochondrial length. Alternatively, direct measurement in mesenchymal cell mitochondria has each cell with approximately 15 mitochondrial connected components, each with approximately 30 branches that are each approximately 1.3 μm long, corresponding to 585 μm total mitochondrial network length (*30*). These two estimates provide a range of approximately 200 – 600 μm for total mitochondrial network length. For 0.048 fission events per second, this mitochondrial length range corresponds to 8×10^-4^ to 2.4×10^-3^ fission events per second per μm in mitochondrial length.

### Data Analysis and Statistics

Graphs and statistical analyses were performed in R. Data represent mean ± SD from at least three independent experiments (n values are indicated in figure legends). Statistical significance was determined by unpaired Student’s t-test (two groups) or one-way ANOVA followed by Tukey’s post hoc test (multiple comparisons).

## Figure Legends

**Supplementary Figure 1.**
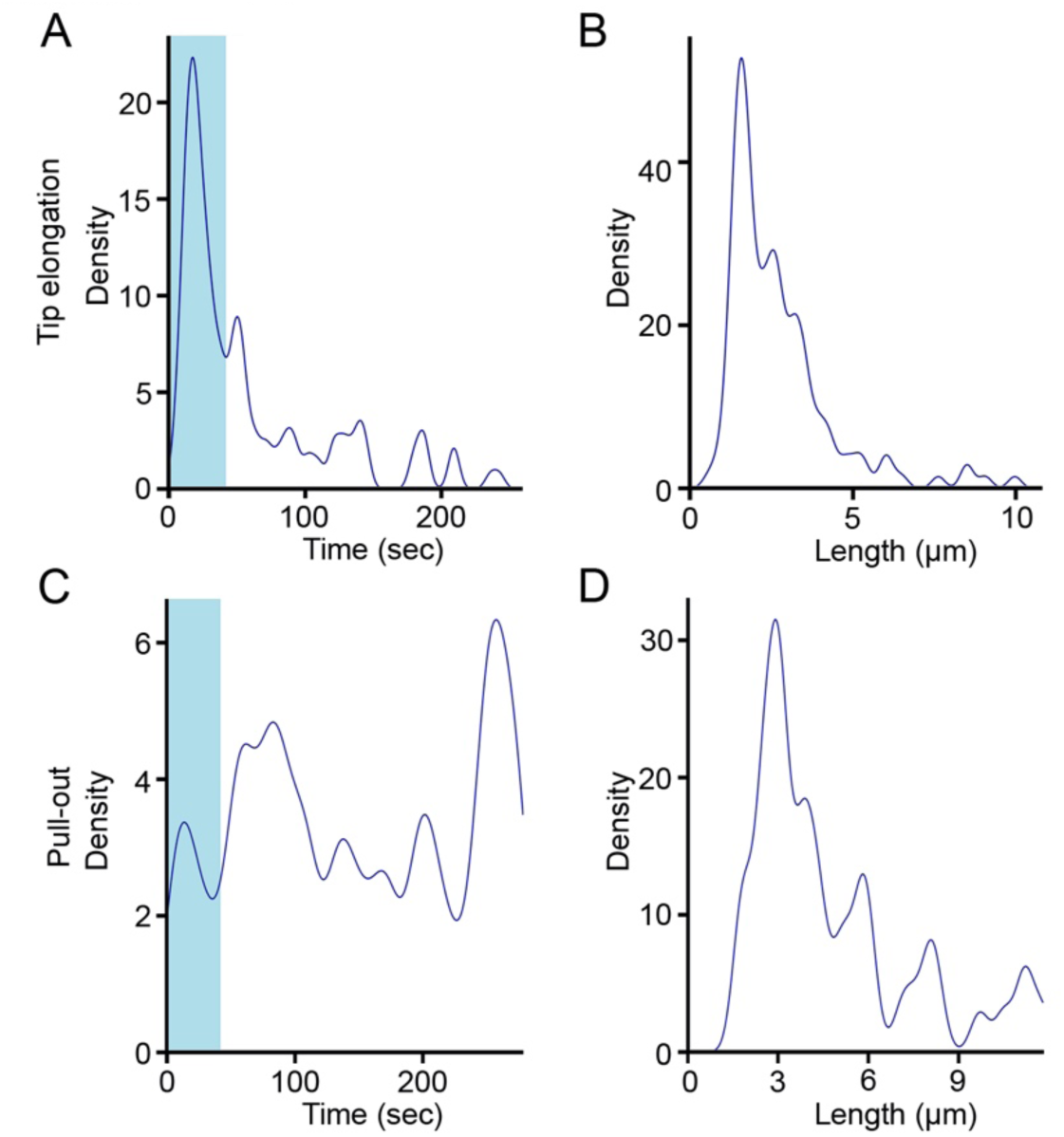
Duration and length of dynamic tubulation events. (A-B) Kinetics of tip elongation in primary human fibroblasts. The duration (A) and length (B) were quantified from 47 events. (C-D) Kinetics of pull-outs in primary human fibroblasts. The duration (C) and length (D) were quantified from 139 events.

